# PODOCYTURIA mRNAs: EARLIER AND SUPERIOR PREDICTORS OF CARDIOVASCULAR OUTCOMES

**DOI:** 10.1101/050039

**Authors:** Assaad A. Eid, Robert H. Habib, Ahmed Abdel Fattah, Nour Al Jalbout, Kamal F. Badr

## Abstract

Increased filtration of albumin, a result of injury to glomerular endothelial and epithelial cells (podocytes), leads to increased albumin excretion rate (AER) which defines “moderate albuminuria”, a predictor of cardiovascular events (CVD). Since filtered albumin is modified by renal proximal tubule albumin retrieval, we hypothesized that urinary podocyte shedding (podocyturia), as a biomarker of ongoing glomerular microvascular injury, may be a more relevant and earlier predictor of CVD than moderate albuminuria.

Type II diabetic subjects (mean age: 46/60 men/women) with normal AER [<2.26 mg/mmole (20 μg/mg) creatinine] and free of overt cardiovascular disease (CVD) were studied. AER and podocyte-specific proteins (podocin and nephrin) mRNAs were measured at baseline (Visit 1), 3-4 years later (Visit 2) and at 7 years (Visit 3). Development of CVD (acute coronary syndrome, stroke, and/or peripheral vascular disease) was collected as outcome. Seven-year Kaplan-Meier time-to-event (log rank) data were compared among baseline biomarker tercile groups (low, intermediate, high).

All three biomarkers increased significantly between visits. Visit 1AER terciles exhibited similar time to CVD (p=0.127), in contrast with step-wise and substantial increases in CVD events predicted by Visit 1 podocin and nephrin terciles. Considered as continuous factors, the covariate-adjusted hazard ratios (95% confidence intervals) [HR] were highest for podocin mRNA [HR=15.9 (6.1-41.8); p<0.001], intermediate for nephrin mRNA [HR=7.61 (3.75-15.5); p<0•001] and lowest for AER [HR=1.17 (1.01-1.36); p=0•041].

Compared to moderate albuminuria, podocyturia predicts more accurately and at a significantly earlier time point the presence of silent systemic vascular injury ultimately manifesting as overt cardiovascular events.

## Introduction

In diabetic and non-diabetic individuals who develop vascular disease, a large body of high-quality evidence has established a direct and continuous relationship between albumin excretion rates (AER) and adverse renal and cardiovascular outcomes, establishing AER as an independent predictor of cardiovascular events (CVD), worse outcomes, and increased mortality in a wide range of clinical settings (1-6). In fact, Ruggegnenti and colleagues (6) demonstrated, in a large population of initially “normoalbuminuric” diabetics, that increasing AER is associated with a continuous nonlinear relationship with CVD development over a mean follow-up period of 9.2 years. Even minimal increases in measurable albuminuria (e.g. from <1 μg/min to 1-2 μg/min) conferred a significant increase in risk (6). Because it is such a powerful and independent predictor of CVD, “moderate albuminuria” (previously “microalbuminuria”) has been incorporated as a major risk factor in several national and international guidelines for the treatment of hypertension and other vascular diseases, and in the calculation of mortality risk in Type II diabetic patients (7).

The mechanisms linking albumin excretion to cardiovascular risk have been the subject of numerous reviews (8-10). Available evidence places endothelial cell injury as a common event occurring simultaneously in systemic and renal vasculature (Figure S-1, online supplement), but detectable only through its measurable functional consequence at the glomerulus (albuminuria) (8,9). Albuminuria is thus a reflection of endothelial injury and is invariably accompanied by injury to subjacent glomerular epithelial cells (podocytes) across which filtered plasma reaches the proximal tubule. Indeed, podocyte shedding (podocyturia) correlates strongly with albuminuria (11-14).

Urinary albumin excretion rate is determined not only by endothelial/podocyte injury and increased albumin filtration, but also by the proportion of filtered albumin taken up by the proximal tubule (14-17). We therefore hypothesized that vascular injury may well be present for some time before the emergence of moderate albuminuria and sought to examine whether we could detect early glomerular (and presumably systemic) endothelial injury as an increase in urinary podocyte shedding (podocyturia), rather than the later onset of moderate albuminuria. We tested this hypothesis in Type II diabetic individuals who presented initially with “normoalbuminuria” [<2.26 mg/mmol (<20 μg/mg) albumin/creatinine excretion rates] and examined whether differences in podocyte shedding rates among these individuals when they are first seen could identify those who would subsequently develop adverse cardiovascular outcomes (acute coronary syndrome, ACS, stroke, peripheral vascular disease, PVD) when followed for seven years.

## Results

### All Patients

Table 1 summarizes the change of all three urinary biomarkers (albumin, podocin mRNA, nephrin mRNA) sampled at three visits (1, 2 and 3) that spanned a median follow-up period of 7.1 years. Briefly, at Visit 1, all 106 subjects in this diabetic study population were free of any known cardiovascular disease and all had “normal” albuminuria levels [AER <2.26 mg/mmol; median (IQR) = 1.07 (0.74-1.19)]. All three urine biomarkers increased over time in all patients in parallel with a gradual decline in eGFR, consistent with progression of diabetic vascular injury (Figure 1). At Visit 2 (at 3 to 4 years), AER had increased substantially [11.9 (10.2-18.8); P<0.001] and this was followed with an additional marked increase at Visit 3 [98.0 (76.8-109.2); P<0.001] (Table 1). The corresponding urinary podocin mRNA [0.65 (0.4-1.1), 4.3 (3.8-5.8) and 10.2 (8.6-12.2) million copies], and nephrin mRNA [1.8 (1.1-2.5), 3.7 (2.3-4.8) and 8.3 (7.3-10.3) million copies] levels also increased progressively between Visits 1 and 3 (all P<0.001) (Figure 1).

**Figure 1:**
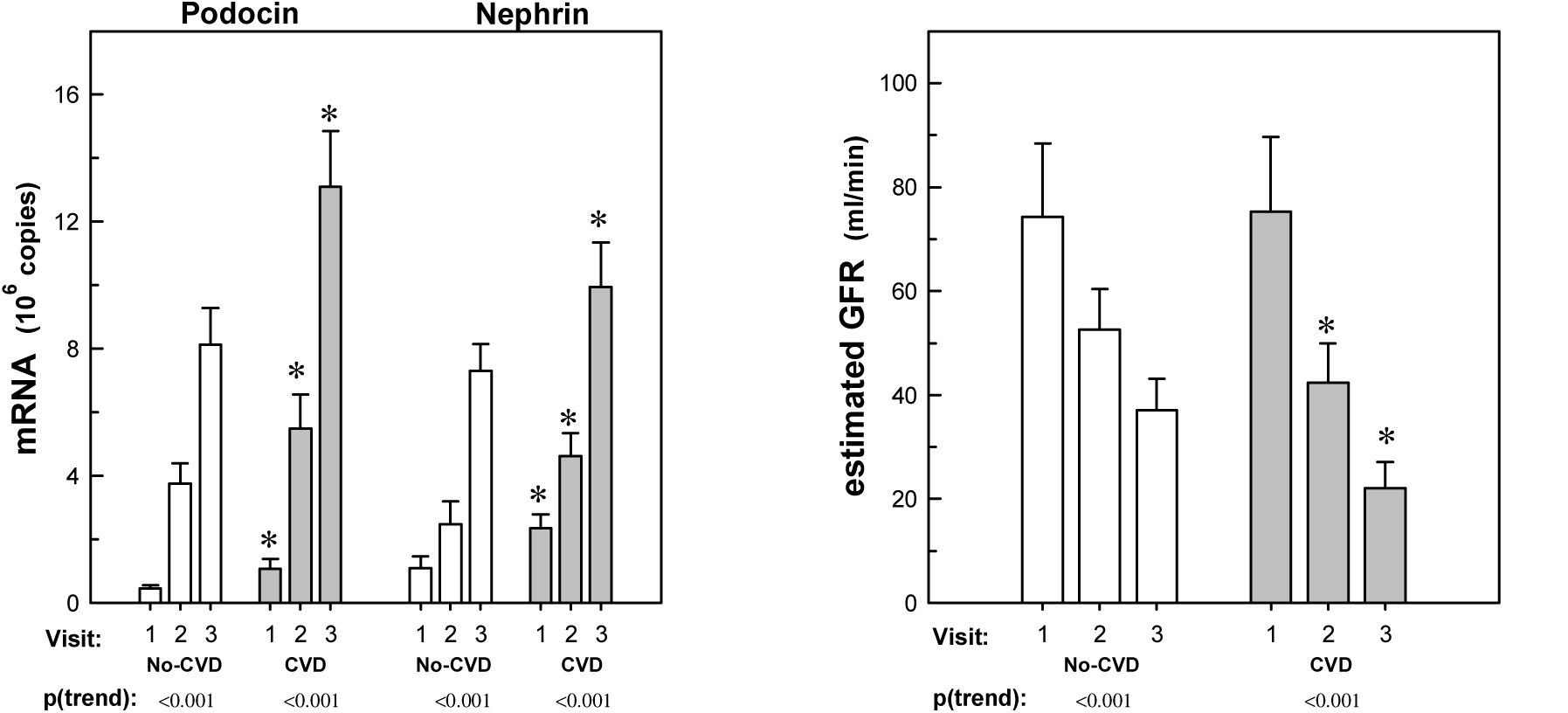
Comparison of mean podocyturia biomarkers (Left) and estimated glomerular filtration rate (eGFR; MDRD formula) measured at visits 1, 2 and 3 for the sub-cohort of diabetic patients that did (CVD) versus did not (No-CVD) develop cardiovascular disease within the 7-year follow-up period. p(trend): within sub-cohort repeated measure analysis of variance analysis testing for significant change with time across visits. *: statistically significantly different compared to same visit No-CVD values.

**Table 1.** Comparison of the rate of increase in urinary biomarkers for Visits 1, 2, and 3.

| Biomarker | Visit 1 |  |  | Visit 2 |  |  | Visit 3 |  |  |
| --- | --- | --- | --- | --- | --- | --- | --- | --- | --- |
|  | Mean±SD | Median | (IQR) | Mean±SD | Median | (IQR) | Mean±SD | Median | (IQR) |
| **<u>All Subjects (N=106)</u>** |  |  |  |  |  |  |  |  |  |
| Podocin (million copies) | 0.78±0.39 | 0.65 | (0.45,1.1) | 4.6±1.3 | 4.3 | (3.8,5.8) | 10.8±2.9 | 10.2 | (8.6,13.2) |
| Nephrin (million copies) | 1.76±0.75 | 1.80 | (1.10,2.50) | 3.6±1.3 | 3.7 | (2.3,4.8) | 8.7±1.8 | 8.3 | (7.3,10.3) |
| AER (mg/mmol Cr) | 1.00±0.26 | 1.07 | (0.74,1.19) | 14.3±4.9 | 11.9 | (10.2,18.8) | 91.0±22.2 | 98 | (76.8,109.2) |
| ΔPodocin (million copies) <sup>#</sup> |  |  |  | 1.05±0.35 | 1.04 | (0.82,1.23) | 1.53±0.60 | 1.45 | (1.07,1.81) |
| ΔNephrin (million copies) <sup>#</sup> |  |  |  | 0.50±0.25 | 0.51 | (0.30,0.66) | 1.05±0.35 | 0.97 | (0.84,1.20) |
| ΔAER (mg/mmol Cr) <sup>#</sup> |  |  |  | 3.62±1.59 | 3.1 | (2.5,4.5) | 13.6±4.5 | 13.7 | (10.4,16.3) |
| **<u>No-CVD (N=50)</u>** |  |  |  |  |  |  |  |  |  |
| Podocin (million copies) | 0.45±0.12 | 0.45 | (0.35,0.54) | 3.7±0.8 | 3.8 | (3.1,4.2) | 8.1±1.2 | 8.4 | (7.3,9.1) |
| Nephrin (million copies) | 1.10±0.36 | 1.10 | (0.90,1.40) | 2.5±0.7 | 2.3 | (1.9,2.8) | 7.3±0.9 | 7.3 | (6.7,8.1) |
| AER (mg/mmol Cr) | 0.96±0.25 | 0.90 | (0.74,1.19) | 11.0±1.4 | 11.10 | (9.95,11.9) | 98.6±14.0 | 99.30 | (89.0,109.2) |
| ΔPodocin (million copies) <sup>#</sup> |  |  |  | 0.88±0.28 | 0.87 | (0.67,1.02) | 1.10±0.24 | 1.07 | (0.94,1.26) |
| ΔNephrin (million copies) <sup>#</sup> |  |  |  | 0.37±0.22 | 0.33 | (0.20,0.51) | 0.89±0.19 | 0.91 | (0.75,1.00) |
| ΔAER (mg/mmol Cr) <sup>#</sup> |  |  |  | 2.71±0.45 | 2.71 | (2.26,3.05) | 13.9±2.8 | 13.70 | (11.5,16.3) |
| **<u>CVD (N=56)</u>** |  |  |  |  |  |  |  |  |  |
| Podocin (million copies) | 1.07±0.31 | 1.10 | (0.80,1.30) | 5.5±1.1 | 5.8 | (4.5,6.4) | 13.1±1.8 | 12.9 | (12.3,14.3) |
| Nephrin (million copies) | 2.36±0.44 | 2.45 | (1.95,2.80) | 4.6±0.7 | 4.8 | (4.1,5.1) | 9.9±1.4 | 10.3 | (8.7,10.9) |
| AER (mg/mmol Cr) | 1.03±0.26 | 1.11 | (0.74,1.26) | 17.1±5.1 | 18.80 | (11.4,20.4) | 84.1±25.7 | 89.50 | (62.7,108.4) |
| ΔPodocin (million copies) <sup>#</sup> |  |  |  | 1.20±0.33 | 1.17 | (1.05,1.33) | 1.91±0.57 | 1.80 | (1.57,2.13) |
| ΔNephrin (million copies) <sup>#</sup> |  |  |  | 0.61±0.22 | 0.61 | (0.49,0.72) | 1.20±0.40 | 1.14 | (0.92,1.32) |
| ΔAER (mg/mmol Cr) <sup>#</sup> |  |  |  | 4.41±1.81 | 4.41 | (3.17,5.43) | 13.2±5.7 | 13.5 | (8.9,16.3) |

All comparisons across visits were significantly different ( $P<0.05$ ). IQR: interquartile range. AER: Albumin excretion rate; <sup>#</sup> Change in biomarker per year relative to Visit 1 value.

### Cardiovascular Disease Sub-cohorts

Urinary biomarker levels in all patients were also analyzed separately for CVD versus No-CVD sub-cohorts during the seven-year follow-up period spanning Visits 1 to 3 (Table 1). Of note, mean baseline (Visit 1) levels of both podocin mRNA (0.45±0.12 vs. 1.07±0.31; P<0.001) and nephrin mRNA (1.10±0.36 vs. 2.36±0.44; P<0.001) in diabetics who did not develop cardiovascular events during the seven-year follow-up period (No-CVD), were significantly lower than those who did develop cardiovascular events (CVD) (Figure 1). Nevertheless, levels in No-CVD individuals were significantly greater than mean values obtained in 20 non-diabetic age-matched volunteers (0.2±0.06 and 0.1±0•02 million copies for nephrin and podocin, respectively). In contrast, AERs at baseline were similar for the No-CVD and CVD subgroups (8.5±2.2 vs. 9.1±2.3; not significant). Comparison of the rate of increase in biomarkers from Visit 1 to Visit 2, and from Visit 2 to Visit 3 revealed a clearly greater rate of rise for all three markers in CVD versus No CVD patients (Table 1 and Figure 1). Tables S-1 and S-2 in the Online Supplement summarize all clinical and biomarker data for CVD and No-CVD patients at baseline and during the subsequent two visits.

### Correlations Among Urinary Biomarkers and Relation to Cardiovascular Disease

Baseline or Visit 1 levels of urinary nephrin and podocin mRNA levels were highly correlated (r2=0.63), but neither was well correlated to the baseline albuminuria levels (r2=0.15 and 0.09 for podocin and nephrin, respectively) (Figure 2-Top). The increases in podocin and nephrin mRNA longitudinally over time appeared to track each other fairly closely and remained well correlated despite the substantial rise in absolute levels (r2=0.40 and 0.37 at Visits 2 and 3, respectively) (Figure 2 Left). Interestingly, the progression from normal to moderate albuminuria observed between Visits 1 and 2 appeared to track with the change in both podocin (r2=0.41) and nephrin (r2=0.34). More importantly, nephrin and podocin mRNA levels were distinctly higher in CVD patients at all visits, including at baseline (Visit 1). This characteristic of urinary podocin and nephrin was however not true for AER which did not discriminate between CVD and No-CVD at baseline (Visit 1). AER was significantly greater for the CVD sub-cohort compared to No-CVD counterparts at Visit 2, but this separation was no longer true at Visit 3.

**Figure 2:**
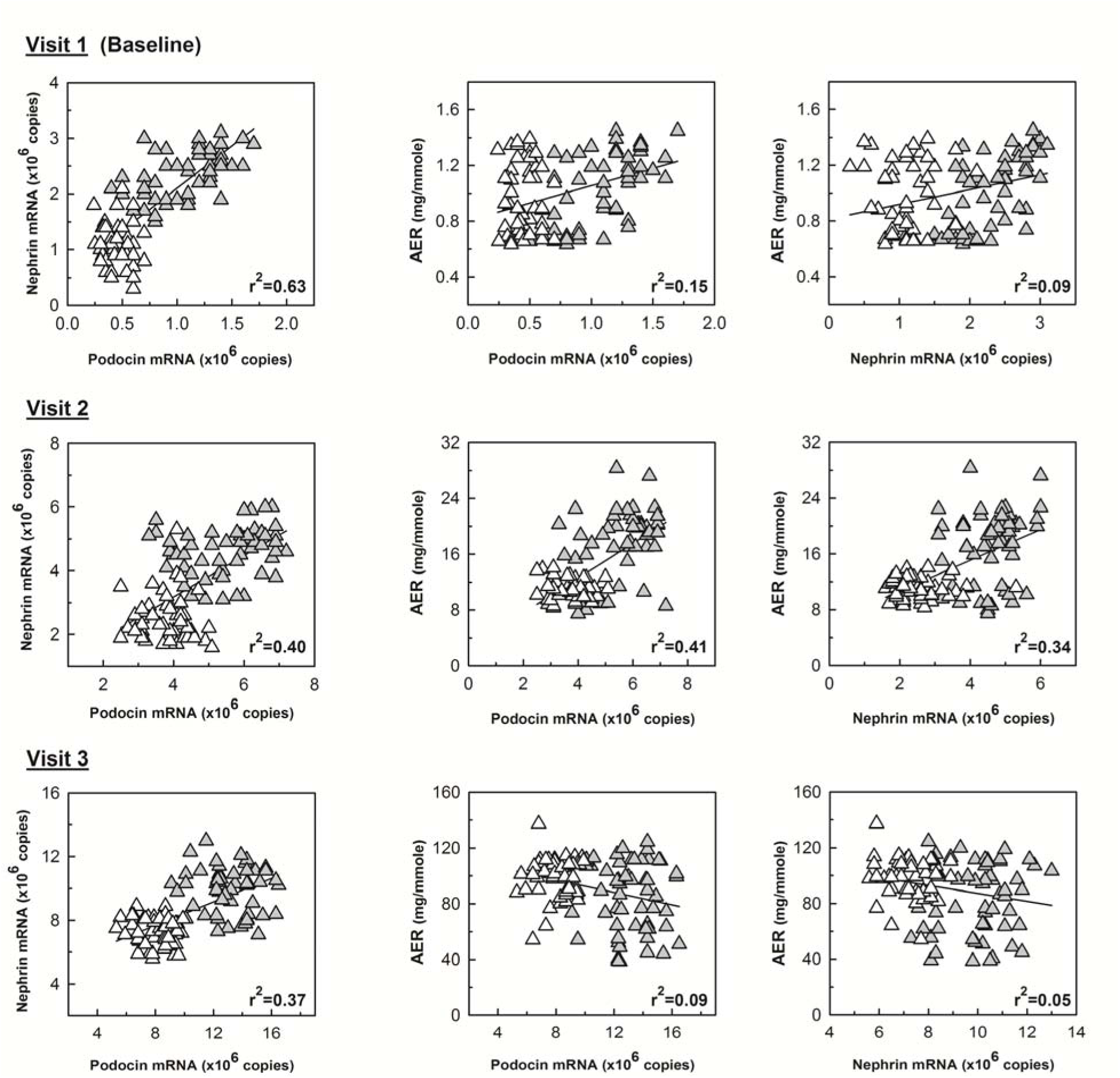
Correlations among podocyturia biomarker levels and albumin excretion rates (AER) shown in individual subjects. Gray: patients developing CVD within 7-year follow-up patients; white: No-CVD subjects; Line: regression line with corresponding correlation coefficient squared (r^2^).

### Time to Cardiovascular Event Analysis

To ascertain the early discriminatory power of each of the urinary biomarkers in predicting CVD, subjects were categorized into 3 sub-groups based on their Visit 1 levels using specific cutoff values for AER (<0.79 (Low); 0.79-1.13 (med) and >1.13 (high) mg/mmol), or using tercile groups in the case of podocin and nephrin mRNA. Comparisons of subject characteristics, clinical data and laboratory values for the podocin and nephrin sub-groups are detailed in Table 2. Development of cardiovascular disease after enrollment was then investigated as a Time-to-Event analysis comparing the derived subgroups for each of the three biomarkers. This analysis indicated that, low, medium and high levels of AER at baseline, albeit all within normal levels (<2.26 mg/mmol), do not separate patients who later developed versus those who did not develop CVD within the 7 year follow-up period (Figure 3-A; Table 3; p=0.127). This was in sharp contrast with the results obtained with either podocin or nephrin mRNA levels (Figure 3-B and 3-C, respectively; Table 3). In the latter cases, increasing levels of podocin and nephrin levels at baseline were highly significantly predictive of development of CVD (both P<0.001), as well as the time to CVD diagnosis or event occurrence. To complement this analysis, we also analysed biomarker levels as continuous variables: the associated covariate-adjusted hazard ratios by multivariate Cox Regression were highest in case of podocin mRNA [HR =15.9 (6.1-41.8); p<0.001], intermediate for nephrin mRNA [HR = 7.61 (3.75-15.5); p<0.001] and lowest for AER [HR=1.17 (1.01-1.36); p=0.041] (Table 3).

**Figure 3:**
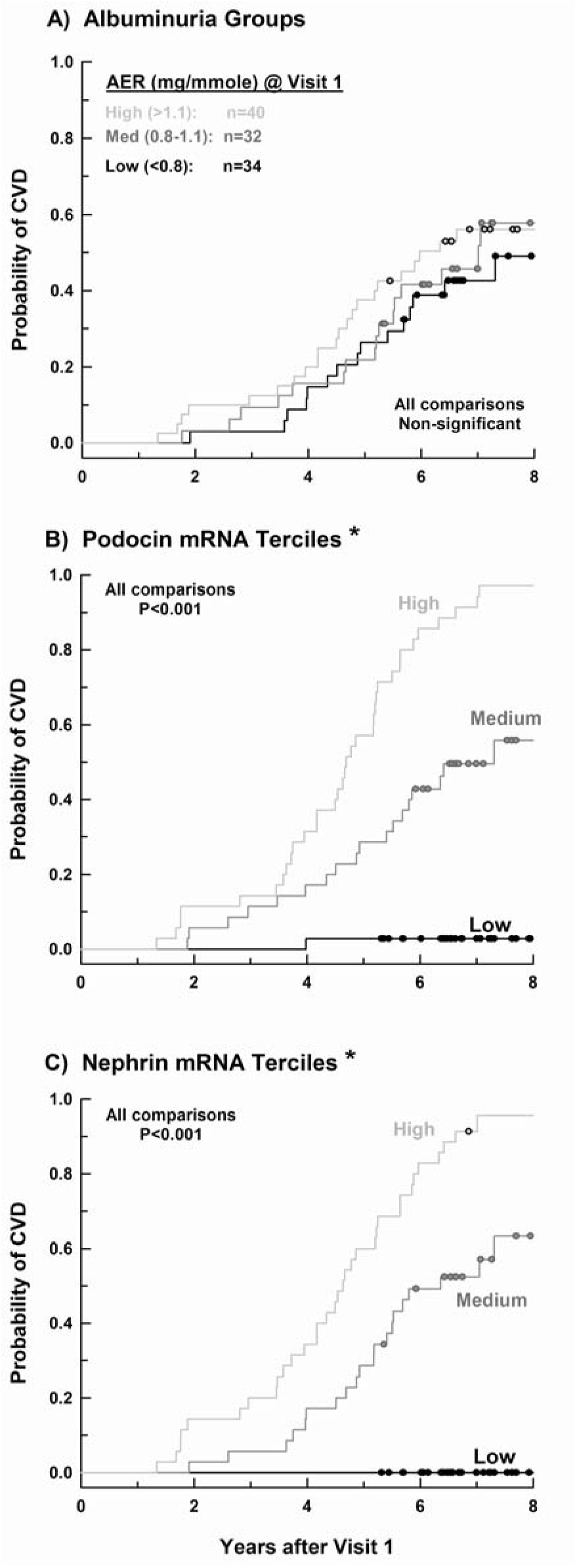
Time to Cardiovascular Event Analysis. Development of cardiovascular disease after patients enrollment as a Time-to-Event analysis for subject sub-cohorts categorized based on levels of: (A) AER, (B) urinary nephrin mRNA and (C) urinary podocin mRNA.

**Table 2:**
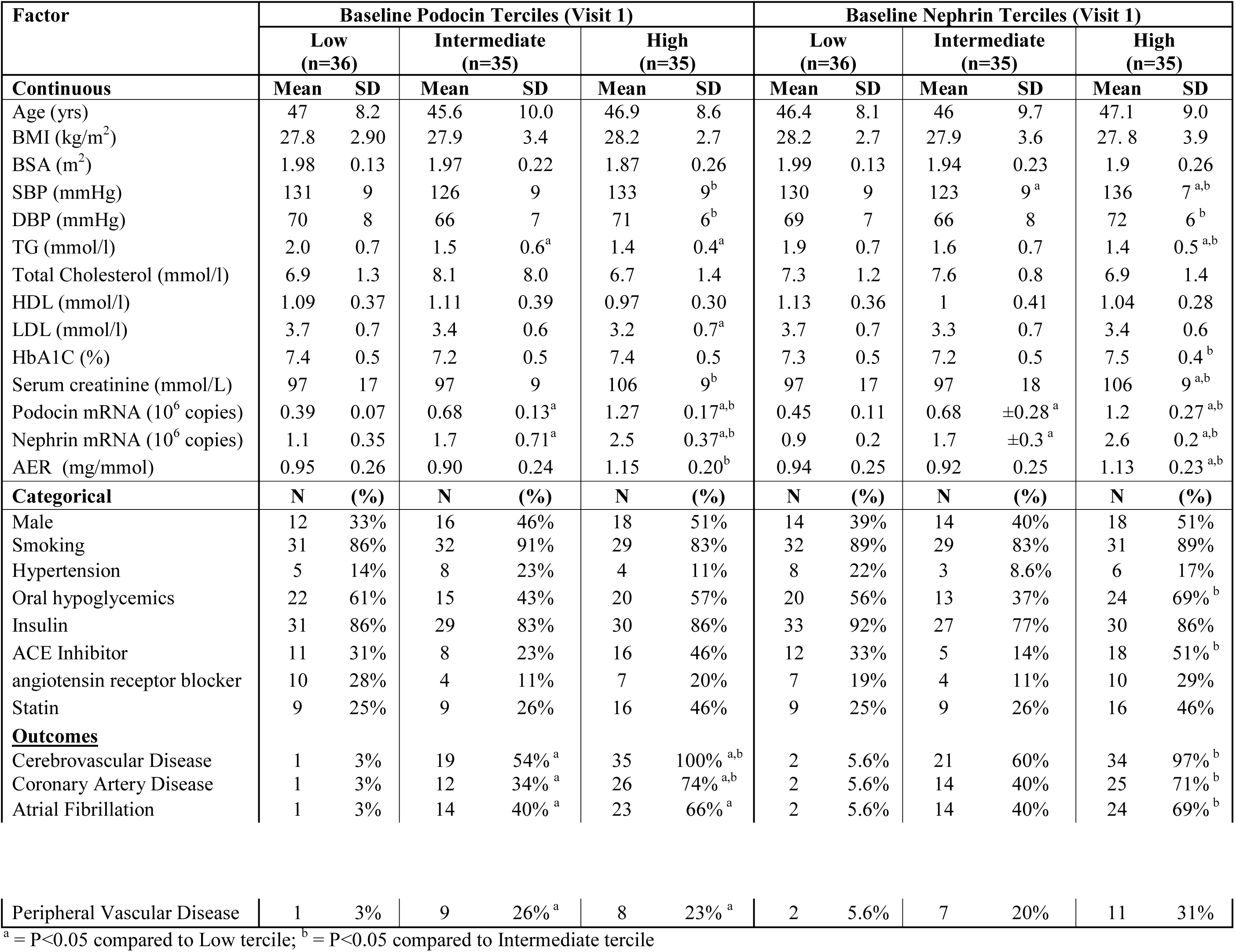
Comparisons of subject characteristics, clinical and laboratory data for baseline podocyturia marker sub-groups

| Factor | Baseline Podocin Terciles (Visit 1) |  |  |  |  |  | Baseline Nephrin Terciles (Visit 1) |  |  |  |  |  |
| --- | --- | --- | --- | --- | --- | --- | --- | --- | --- | --- | --- | --- |
|  | Low<br>(n=36) |  | Intermediate<br>(n=35) |  | High<br>(n=35) |  | Low<br>(n=36) |  | Intermediate<br>(n=35) |  | High<br>(n=35) |  |
| Continuous | Mean | SD | Mean | SD | Mean | SD | Mean | SD | Mean | SD | Mean | SD |
| Age (yrs) | 47 | 8.2 | 45.6 | 10.0 | 46.9 | 8.6 | 46.4 | 8.1 | 46 | 9.7 | 47.1 | 9.0 |
| BMI (kg/m<sup>2</sup>) | 27.8 | 2.90 | 27.9 | 3.4 | 28.2 | 2.7 | 28.2 | 2.7 | 27.9 | 3.6 | 27.8 | 3.9 |
| BSA (m<sup>2</sup>) | 1.98 | 0.13 | 1.97 | 0.22 | 1.87 | 0.26 | 1.99 | 0.13 | 1.94 | 0.23 | 1.9 | 0.26 |
| SBP (mmHg) | 131 | 9 | 126 | 9 | 133 | 9<sup>b</sup> | 130 | 9 | 123 | 9<sup>a</sup> | 136 | 7<sup>a,b</sup> |
| DBP (mmHg) | 70 | 8 | 66 | 7 | 71 | 6<sup>b</sup> | 69 | 7 | 66 | 8 | 72 | 6<sup>b</sup> |
| TG (mmol/l) | 2.0 | 0.7 | 1.5 | 0.6<sup>a</sup> | 1.4 | 0.4<sup>a</sup> | 1.9 | 0.7 | 1.6 | 0.7 | 1.4 | 0.5<sup>a,b</sup> |
| Total Cholesterol (mmol/l) | 6.9 | 1.3 | 8.1 | 8.0 | 6.7 | 1.4 | 7.3 | 1.2 | 7.6 | 0.8 | 6.9 | 1.4 |
| HDL (mmol/l) | 1.09 | 0.37 | 1.11 | 0.39 | 0.97 | 0.30 | 1.13 | 0.36 | 1 | 0.41 | 1.04 | 0.28 |
| LDL (mmol/l) | 3.7 | 0.7 | 3.4 | 0.6 | 3.2 | 0.7<sup>a</sup> | 3.7 | 0.7 | 3.3 | 0.7 | 3.4 | 0.6 |
| HbA1C (%) | 7.4 | 0.5 | 7.2 | 0.5 | 7.4 | 0.5 | 7.3 | 0.5 | 7.2 | 0.5 | 7.5 | 0.4<sup>b</sup> |
| Serum creatinine (mmol/L) | 97 | 17 | 97 | 9 | 106 | 9<sup>b</sup> | 97 | 17 | 97 | 18 | 106 | 9<sup>a,b</sup> |
| Podocin mRNA (10<sup>6</sup> copies) | 0.39 | 0.07 | 0.68 | 0.13<sup>a</sup> | 1.27 | 0.17<sup>a,b</sup> | 0.45 | 0.11 | 0.68 | ±0.28<sup>a</sup> | 1.2 | 0.27<sup>a,b</sup> |
| Nephrin mRNA (10<sup>6</sup> copies) | 1.1 | 0.35 | 1.7 | 0.71<sup>a</sup> | 2.5 | 0.37<sup>a,b</sup> | 0.9 | 0.2 | 1.7 | ±0.3<sup>a</sup> | 2.6 | 0.2<sup>a,b</sup> |
| AER (mg/mmol) | 0.95 | 0.26 | 0.90 | 0.24 | 1.15 | 0.20<sup>b</sup> | 0.94 | 0.25 | 0.92 | 0.25 | 1.13 | 0.23<sup>a,b</sup> |
| Categorical | N | (%) | N | (%) | N | (%) | N | (%) | N | (%) | N | (%) |
| Male | 12 | 33% | 16 | 46% | 18 | 51% | 14 | 39% | 14 | 40% | 18 | 51% |
| Smoking | 31 | 86% | 32 | 91% | 29 | 83% | 32 | 89% | 29 | 83% | 31 | 89% |
| Hypertension | 5 | 14% | 8 | 23% | 4 | 11% | 8 | 22% | 3 | 8.6% | 6 | 17% |
| Oral hypoglycemics | 22 | 61% | 15 | 43% | 20 | 57% | 20 | 56% | 13 | 37% | 24 | 69%<sup>b</sup> |
| Insulin | 31 | 86% | 29 | 83% | 30 | 86% | 33 | 92% | 27 | 77% | 30 | 86% |
| ACE Inhibitor | 11 | 31% | 8 | 23% | 16 | 46% | 12 | 33% | 5 | 14% | 18 | 51%<sup>b</sup> |
| angiotensin receptor blocker | 10 | 28% | 4 | 11% | 7 | 20% | 7 | 19% | 4 | 11% | 10 | 29% |
| Statin | 9 | 25% | 9 | 26% | 16 | 46% | 9 | 25% | 9 | 26% | 16 | 46% |
| **Outcomes** |  |  |  |  |  |  |  |  |  |  |  |  |
| Cerebrovascular Disease | 1 | 3% | 19 | 54%<sup>a</sup> | 35 | 100%<sup>a,b</sup> | 2 | 5.6% | 21 | 60% | 34 | 97%<sup>b</sup> |
| Coronary Artery Disease | 1 | 3% | 12 | 34%<sup>a</sup> | 26 | 74%<sup>a,b</sup> | 2 | 5.6% | 14 | 40% | 25 | 71%<sup>b</sup> |
| Atrial Fibrillation | 1 | 3% | 14 | 40%<sup>a</sup> | 23 | 66%<sup>a</sup> | 2 | 5.6% | 14 | 40% | 24 | 69%<sup>b</sup> |
| Peripheral Vascular Disease | 1 | 3% | 9 | 26%<sup>a</sup> | 8 | 23%<sup>a</sup> | 2 | 5.6% | 7 | 20% | 11 | 31% |

<sup>a</sup> = P<0.05 compared to Low tercile; <sup>b</sup> = P<0.05 compared to Intermediate tercile

**Table 3.** Hazard Ratios for the development of cardiovascular disease derived for all three urinary biomarkers considered as continuous or categorical biomarkers.

| **Biomarker** | **N** | **B** | **SE** | **p value** | **Adjusted HR**<br>**(95% CI)** |
| --- | --- | --- | --- | --- | --- |
| **<u>Continuous Values</u>** |  |  |  |  |  |
| AER (mcg/mg) | 106 | .155 | .076 | .041 | 1.17 (1.01-1.36) |
| Podocin mRNA (10<sup>6</sup> copies) | 106 | 2.767 | .492 | .000 | 15.9 (6.1-41.8) |
| Nephrin mRNA (10<sup>6</sup> copies) | 106 | 2.030 | .361 | .000 | 7.61 (3.75-15.5) |
| **<u>Two Categories<sup>#</sup></u>** |  |  |  |  |  |
| AER | 52/54 | .502 | .329 | .127 | 1.65 (0.87-3.14) |
| Podocin mRNA | 53/53 | 2.507 | .506 | .000 | 12.3 (4.55-33.1) |
| Nephrin mRNA | 52/54 | 2.767 | .520 | .000 | 15.9 (5.74-44.0) |
| **<u>Three Categories</u>** |  |  |  |  |  |
| AER |  |  |  | .289 |  |
| Low | 34 |  |  |  | 1.0 (ref) |
| Intermediate | 32 | .426 | .430 | .322 | 1.53 (0.66-3.55) |
| High | 40 | .610 | .390 | .118 | 1.84 (0.86-3.95) |
| Podocin terciles |  |  |  | .000 |  |
| Low | 36 |  |  | .000 | 1.0 (ref) |
| Intermediate | 35 | 3.003 | 1.049 | .004 | 20.1 (2.58-157.4) |
| High | 35 | 4.086 | 1.039 | .000 | 59.5 (7.8-456.2) |
| Nephrin terciles * |  | * | * | * | * |

N = number of subjects; B (SE) = cox regression coefficient (standard error); HR (95% CI) = hazard ratio (95% confidence interval). Data adjusted for the following Visit 1 factors: gender, age, body mass index, smoking, hypertension (yes/no), statin use (yes/no), systolic and diastolic blood pressure, triglyceride, low and high density lipoprotein, and hemoglobin A1c levels.

## Discussion

Under physiologic (“normal”) conditions, an appreciable amount of albumin is filtered across glomerular epithelial cells (podocytes) [15-20]. In healthy children and adults, the amount of albumin appearing in the urine is the algebraic difference between filtered albumin and the amount retrieved by the S1 segment of the proximal tubule [15,16,19,20]. An increasingly large body of evidence in humans and other mammals assigns a larger role for proximal tubule albumin retrieval in determining urinary albumin excretion than previously appreciated [15,16,19,20]. Studies have established that proximal tubule albumin retrieval rates are highly regulated and are also influenced by total proximal tubule mass and disease states, including diabetes [16,19,20]. It therefore seems reasonable to assume that individuals vary widely in the proximal handling of filtered loads of albumin and fractional albumin retrieval. The tight coupling between AER and CV outcomes, however, suggests strongly that it is progressive glomerular and systemic vascular injury, leading to an increase in filtered albumin, that underlies progressive albuminuria in these individuals. For urinary albumin to reflect filtered albumin, however, proximal tubule retrieval of albumin must be saturated. Only then would vascular injury (filtered albumin) correlate positively and tightly with urinary albumin, a relationship repeatedly demonstrated in numerous large population studies [1,2,4,5,6,21,22]. In an individual patient, however, the time elapsing between the onset of vascular injury and the saturation of proximal retrieval cannot be known; increased filtration of albumin may be present for years before urinary albumin levels begin to rise.

Our study addressed this hypothesis by targeting an earlier event in the vascular injury pathway, podocyte shedding, and assessing the capacity of podocyturia to predict CV outcomes before AER increases above the “normal” range. Podocyturia correlates strongly with proteinuria and renal functional deterioration in diabetic and non-diabetic individuals, as well as in patients with inflammatory and non-inflammatory glomerular diseases [23-27], but it has not been previously evaluated as an early predictor of systemic vascular injury. Podocyturia is best measured by realtime PCR quantitation of mRNA for unique podocyte-specific proteins (such as nephrin, podocalyxin, podocin, synaptopodin, and others) [28,29].

Our study population consisted of relatively young diabetics (mean age 47 years at Visit 1) who, despite being “normoalbuminuric”, carried other risk factors for progression of CVD, including a high proportion of smokers and overweight/obese individuals, coupled to relatively poor glycemic control (Table 2 and Table S-1 in Online Supplement). Not surprisingly, therefore, there were progressive increases in all three urinary biomarkers (AER and nephrin/podocin mRNA) between visits 1, 2 and 3, reflecting progressive vascular injury. Despite normal baseline AER levels, more than half of the subjects (56/106) developed cardiovascular disease within the 7 year follow-up period. Importantly, however, those that did versus did not develop CVD had similar baseline AER at enrollment, but significantly different podocin and nephrin mRNA (Figure 1, Table 1). The rate of increase in biomarkers was accelerated in CVD versus No-CVD patients, supporting the notion that podocyte injury and albuminuria are indeed a reflection of progressive endothelial/vascular injury occurring silently and simultaneously in the cerebral, coronary, and systemic circulation.

The data summarized in Figure 2 provides evidence for the superiority of nephrin/podocin mRNA over AER as a faithful marker of CV injury over time. Nephrin and podocin mRNA levels correlated well with each other, even as injury progressed over seven years, suggesting that podocyte shedding is indeed well reflected by both measurements. More importantly, at all three visits, CVD patients had distinctly higher values of urinary podocin/nephrin mRNA than their No-CVD counterparts. That AER at Visit 1 did not correlate with nephrin or podocin mRNA, nor did it distinguish between CVD and No-CVD patients, supports strongly the notion that the major determinant of AER at this early stage of injury is not filtered albumin, but the fraction of it retrieved in the proximal tubule, as suggested by others [15,16,19,20]. At Visit 2, AER had increased to levels within the moderate albuminuria range (previously termed “microalbuminuria”) and indeed at this stage of injury, at which time filtered albumin contributes increasingly to the final value of AER, the well–established capacity of moderate albuminuria to distinguish CVD from No-CVD is evident and AER correlates well with both nephrin and podocin mRNA levels. By Visit 3, however, AER had increased to the macroalbuminuric range and is no longer relevant to identifying CV risk. Even at Visit 3, however, urinary podocin/nephrin mRNAs continues to distinguish the two patient groups (Bottom Panel of Figure1).

## Limitations

Despite the promising and provocative findings, our study has certain limitations that may reflect on its generalizability. Specifically, our study population consisted of a relatively small number of Type II diabetic patients who had normo-albuminuria at enrollment. Arguably, the applicability of the claimed superiority of podocyturia compared to AER to non–-diabetics and the elderly may then be challenged. We could not address the elderly issue in the present analysis, which should be addressed in a future age–stratified analysis. Of note, normo-albuminuria in elderly diabetics is not common. Our choice of the diabetic population was in fact purposeful, since we expected accelerated endothelial injury compared to non–diabetics, and hence a relatively more frequent and earlier onset of CVD within 7 years. It is important to note that our study was meant to be a proof of concept proposing novel biomarkers of vascular injury, and hence CVD, as opposed to being an exhaustive assessment of these biomarkers. The dramatic statistical differences observed even with this small number of patients provides compelling evidence for the underlying hypothesis. We expect that the correlation between podocyturia and systemic vascular injury will be present in non–-diabetics as well. This too awaits future studies.

In conclusion, our study highlights the potential usefulness of urinary podocyte mRNA quantitation as a surrogate marker for systemic silent microvascular injury. The principal benefit of this assay would be its capacity to identify individuals destined to develop moderate albuminuria and vascular disease several years before either of these outcomes is detectable by present methodologies. Compared to AER, podocyturia appears to be a more accurate biomarker for the presence of preclinical (silent) systemic vascular injury subsequently manifesting as overt cardiovascular events. In addition, it is capable of capturing at-risk individuals several years earlier than albuminuria, providing a valuable opportunity for preventive interventions. We contend that, pending standardization of urinary podocyte mRNA measurements and independent verification in other cohorts, prospective studies, and other disease states, a clinically applicable assay for podocyte-specific mRNAs will potentially replace AER as the best predictor of systemic vascular injury. Should this be the case, it will be due to the remarkable inherent capacity of glomerular endothelial cells to “report” (through podocyturia) on the health of the systemic endothelium. “Amplification” of the injury “signal” by the large amounts of filtered fluid and the capacity to detect injury markers in the urine, suggest a novel role for the renal glomerulus as an endogenous “endotheliometer”, providing crucial information on the presence, severity and progression of pre-clinical systemic vascular injury.

## Methods

### Study design and Patients

This is a clinical cohort follow-up study in a retrospective patient series participating in a longitudinal biologic sample repository, with additional biomarker measurements. Enrolled patients were either outpatients or in-patients with Type II diabetes mellitus seen for initial presentation and then followed between 1999 and 2014. A single leftover urine sample from age-matched twenty healthy (non-diabetic and no cardiovascular disease) controls was also collected for measurement of baseline healthy biomarkers levels.

### Population

From the large cohort of diabetic patients enrolled for biomarker studies at Hamad General Hospital, Doha – Qatar, we identified and preselected 106 diagnosed Type 2 diabetic patients (age: 25–89 years; men/women: 46/60) on whom follow up visits (clinical data and urine samples) were available and complete for seven years since their initial visit, whose urinary albumin excretion rates on Visit 1 were below what was traditionally termed “microalbuminuria” [<2.26 mg/mmol (<20 microgram/mg) albumin/creatinine], and who had no history of cardiovascular diseases of any kind, in particular ACS, stroke or PVD. Patients were pre-selected to have survived for 7 years since Visit 1. Hypertension (present in 17 patients on Visit 1) was not considered an adverse CV outcome.

### Study approval

The institutional research board of Hamad General Hospital, Doha - Qatar, approved this study (and consent was secured from study subjects to use their left over urine samples and to review their medical charts for blood/urine test results and for all relevant clinical diagnoses (Subject Ethical Approval Research Protocol #11313/11).

### Sample Collection and Storage

Urine and blood samples were collected during Visit 1 along with a complete medical history and physical examination. Subjects were generally seen at 6 months intervals. Follow-up visits were reviewed for approximately 7 years after Visit 1 to obtain interval history, physical examination, and blood and urine test results at each Visit. Clinical data, including new onset CVD, as well as urine and blood samples were analyzed from Visit 1, Visit 2 (mean follow up after Visit 1 of 3.8 years; SD 0.6 years) and Visit 3 (mean follow up after Visit 2 of 3.2 +/- 0.7 years).

After collection, urine sample are kept at 4°C (for a maximum of 24 hours) until moved to the lab where they are kept at −80°C for prolonged storage period. Urine RNA stability is in direct correlation with the temperature of storage of the urine. The influence of storage temperature and freeze–thaw cycles on urinary RNA stability and integrity was examined at temperatures ranging from room temperature to −80°C over a period of 20 days; −70°C proved the optimal temperature in terms of future RNA sample stability, since the RNA samples were degraded by almost 50 to 60 % when kept between 4°C and −10°C. RNA degradation was also detected in samples stored at −20°C and −40°C; no such effect were detected with storage at −80°C. Urine samples were analyzed for albumin excretion rates (AER; mg/mmol creatinine) and for podocin and nephrin mRNA levels (see below). All samples were assigned a study number that linked them to the clinical information in clinical charts. Samples were coded and analyzed with the lab operator blinded to the Visit number, the subject it belonged to, or whether this subject eventually developed CVD or not.

## Procedures

### Volume of Urine for Assay

We assumed that the samples contain low mRNA levels. Podocin and nephrin mRNA were not detectable in 25 to 30% of samples starting with < 1 ml of urine. Therefore, 1.7 ml were used in all assays.

### Urine Processing

Urine was centrifuged at 4°C for 15 minutes at 4000 rpm (3200 g) on a table-top centrifuge. The low-speed centrifugation conditions used do not pellet exosomes and other subcellular structures, so we assumed that the urine pellet mRNAs measured are derived from whole cells and are therefore a measure of the podocyte detachment rate. The supernatant was removed and stored at −20°C for protein, creatinine, and other measurements. The urine pellet was suspended in 1.5 ml of cold diethylpyrocarbonate-treated PBS (pH, 7.4) at 4°C. The pellet material in 1.5 ml PBS was then centrifuged at 12,000 rpm in a mini-centrifuge for 5 minutes at 4°C. The supernatant was discarded. 350 μl of RLT buffer was added to the washed pellet, containing β-mercaptoethanol at 10 μl/ml of RLT buffer, according to the manufacturer protocol (RNeasy kit; Qiagen - Germantown, MD). The pellet was suspended in RLT/β-mercaptoethanol buffer and then frozen at −80°C for assay.

### RNA Preparation and Quantitative RT-PCR Assay

The total urine pellet RNA was isolated using the manufacturer protocol (RNeasy Mini Kit; Qiagen - Germantown, MD). Quantitation of the absolute nephrin and podocin, mRNA abundance was performed using the CFX96 Touch™ Real-Time PCR Detection System (Bio-rad, USA) using TaqMan Fast Universal PCR Master Mix, with sample cDNA in a final volume of 25 μl per reaction. TaqMan Probes (Applied Biosystems) used were as follows: human NPHS1 (nephrin gene accession number: Q1KMS5) and NPHS2 (podocin gene accession number: Q9NP85). All data were from 2-μl sample measured in duplicate. Standard curves were constructed for each assay using serially diluted cDNA standards. Assays were accepted only if the r2 > 0.97 for standard curves. Human nephrin and podocin cDNA of known sequence and concentration were used as standards for each assay so that the data could be calculated on number of copy basis for each probe. RNA urine analysis quality, recovery, and stability were performed.

## Statistics

Continuous variables are summarized as mean±standard deviation or median (inter-quartile range) as applicable. Categorical variables were summarized as counts and percentages. Study subjects were grouped into tercile sub-groups based on their baseline (Visit 1) AER, nephrin, and podocin mRNA levels (low, intermediate and high). Baseline characteristics were compared among the three groups. The primary outcome of interest was the time for diagnosis of CVD, which included documentation of any of the following during the period of follow up for each patient: coronary artery disease, myocardial infarction, heart failure, cerebrovascular accident (stroke) and PVD. Time-to-Event (CVD diagnosis) analysis was performed via Product-Limit or Kaplan-Meier estimates, which were compared across terciles. Cox regression was used to adjust for confounders (SPSS 21; IBM, NY). Rate of rise in nephrin and podocin urinary mRNA, as well as albuminuria levels, was compared across study groups via repeated measures of analysis of variance analysis (RM-ANOVA). A p value less than 0.05 was used to indicate statistical significance.

## Acknowledgement

AAE received funding from the Juvenile Diabetes Research Foundation and the Medical Practice Plan of the American University of Beirut Medical Center (AUBMC). KFB received funding from the Dean’s Support Package to the “Vascular Medicine Program” at AUBMC.

### Contributions

KFB conceived and designed the study. AFHA and AAE recruited subjects and collected the data. AAE performed the mRNA studies. NA performed data management, literature review and reference management. RHH performed data analysis. RHH, KFB and AAE interpreted the findings. KFB wrote the first draft. All authors drafted, reviewed, and completed and approved the manuscript and the decision to submit for publication.

